# Inhibition of LRRK2 kinase activity rescues deficits in striatal dopamine dynamics in VPS35 p.D620N knock-in mice

**DOI:** 10.1101/2023.05.14.540654

**Authors:** Mengfei Bu, Jordan Follett, Igor Tatarnikov, Shannon Wall, Dylan Guenther, Isaac Deng, Genevieve Wimsatt, Austen Milnerwood, Mark S. Moehle, Habibeh Khoshbouei, Matthew J. Farrer

**Affiliations:** Department of Neurology and Norman Fixel Institute for Neurological Diseases, University of Florida, Gainesville, FL, USA; Djavad Mowafaghian Centre for Brain Health, University of British Columbia, Vancouver, Canada; Montreal Neurological Institute-Hospital, McGill University, Montreal, Canada; Department of Pharmacology and Therapeutics, University of Florida, Gainesville, FL, USA; Department of Neuroscience, University of Florida, Gainesville, FL, USA

## Abstract

Dysregulation of dopamine neurotransmission profoundly affects motor, motivation and learning behaviors, and is often observed during the prodromal phase of Parkinson’s disease (PD). However, the mechanism underlying these pathophysiological changes remains to be elucidated. Mutations in vacuolar protein sorting 35 (*VPS35*) and leucine-rich repeat kinase 2 (*LRRK2*) both lead to autosomal dominant PD, and VPS35 and LRRK2 may physically interact to govern the trafficking of synaptic cargos within the endo-lysosomal network in a kinase-dependent manner. To better understand the functional role of VPS35 and LRRK2 on dopamine physiology, we examined Vps35 haploinsufficient (Haplo) and Vps35 p.D620N knock-in (VKI) mice and how their behavior, dopamine kinetics and biochemistry are influenced by LRRK2 kinase inhibitors. We found Vps35 p.D620N significantly elevates LRRK2-mediated phosphorylation of Rab10, Rab12 and Rab29. In contrast, Vps35 haploinsufficiency reduces phosphorylation of Rab12. While striatal dopamine transporter (DAT) expression and function is similarly impaired in both VKI and Haplo mice, that physiology is normalized in VKI by treatment with the LRRK2 kinase inhibitor, MLi-2. As a corollary, VKI animals show a significant increase in amphetamine induced hyperlocomotion, compared to Haplo mice, that is also abolished by MLi-2. Taken together, these data show Vps35 p.D620N confers a gain-of-function with respect to LRRK2 kinase activation, and VPS35 and LRRK2 functionally interact to regulate DAT trafficking and striatal dopamine neurotransmission.

## Introduction

Parkinson’s disease (PD) is the most common aged-related movement disorder. Motor symptoms are associated with the insidious but progressive loss of striatal dopaminergic innervation that ascends from neuronal soma in the midbrain. Most patients with late-onset PD are idiopathic and have a multifactorial etiology. This includes a well-defined background of polygenic risk, albeit of marginal effect^1^, and rare patients have a monogenic component^2^. Dominantly-inherited mutations in leucine-rich repeat kinase 2 (*LRRK2* p.N1437H, p.R1441C/G/H, p.Y1699C, p.G2019S and p.I2020T)^3–6^ and vacuolar protein sorting 35 (*VPS35* p.D620N)^7, 8^ have been genetically linked to parkinsonism that is clinically and pathologically indistinguishable from idiopathic PD^2^. These mutations are some of the most penetrant causes of PD, and confer high genotypic risk.

VPS35 is an essential component of the retromer complex required for endosomal sorting and recycling of cargo proteins, that precludes their endo-lysosomal degradation^9^. *In vitro* studies of Vps35 p.D620N have shown that the mutation does not affect protein expression nor disturbs retromer complex assembly^10^. However, many studies have found the amino acid substitution compromises cargo sorting functions of the retromer complex, consistent with a partial loss-of-function^10, 11^. VPS35 deficiency also impairs mitochondrial fusion and induces age-dependent α-synuclein pathology^12–14^. Previously we showed Vps35 p.D620N knock-in mice (VKI) have early alterations within the dopaminergic system in the absence of any overt motor phenotypes^15^. Subsequent studies have demonstrated that VKI mice exhibit age-dependent neurodegeneration and tau pathology^16, 17^. However, the pathophysiological role of VPS35 in the dopaminergic system is unknown, and there are conflicting reports regarding the p.D620N mutation as it is unclear whether it’s a gain or loss of function^11^.

Pathogenic *LRRK2* mutations constitutively activate LRRK2 kinase activity, directly^18^ and indirectly^19–21^. Brain imaging in clinically asymptomatic *LRRK2* heterozygotes reveals physiological differences in dopaminergic and serotonergic systems^22^. LRRK2 kinase activity is also significantly elevated in neutrophils and monocytes from heterozygous patients with *VPS35* p.D620N parkinsonism^23^, and similarly observed in embryonic fibroblasts and whole tissue obtained from VKI mice^23^. Multiple studies have shown that VPS35 and LRRK2 physically and functionally interact to regulate vesicular trafficking^23–25^. Overexpression of *Vps35* may rescue locomotor deficits in Drosophila expressing mutant *Lrrk2*^26^ but the strict 1:1:1 stoichiometry of VPS35:VPS29:VPS26 subunits (the retromer ‘core’ trimer) suggests results from non-physiologic overexpression must be interpreted cautiously. Simple overexpression of VPS35 is toxic in primary neuronal culture and leads to a 50% reduction in synapses^27^. Nevertheless, physiologic interactions between LRRK2 and VPS35 remain to be explored given the therapeutic potential of these targets.

To assess gain-versus loss-of-function of Vps35 p.D620N, and to better understand the relationship between VPS35 and LRRK2, we performed a parallel comparison of Vps35 p.D620N (VKI) and Vps35 haploinsufficient (Haplo) mice (in which Vps35 expression is reduced by 50%). Outcome measures focused on LRRK2 biology, and the interplay between LRRK2 and VPS35 in the dopamine system. Here, we demonstrate that both VKI and Haplo animals exhibit a loss-of-function in DAT trafficking and dopamine reuptake. Nevertheless, their behaviour is distinct as amphetamine-induces comparable hyperlocomotion in VKI and wildtype mice, but not Haplo mice. Furthermore, a priori LRRK2 kinase inhibition ablates amphetamine-induced hyperlocomotion in VKI animals and normalizes DAT expression and function.

## Methods

### Animals

Vps35 constitutive knock-in (VKI) and haploinsufficient mice (Haplo) were concomitantly generated by Ozgene PLC (Australia), as previously described^15^. A floxed “mini-gene” consisting of: (1) splice acceptor, Vps35 exon 15–17 coding sequence, a polyadenylation signal (pA), and; (2) a PGK-neomycin -pA selection cassette (neo) internally flanked by FRT sites, was inserted into intron 14. The 5′ targeting arm spanning endogenous exon 15 was used to introduce the Vps35 g.85,263,520G>A (p.D620N) mutation (GRCm38/mm10; NM_022997.5). In this design, the mini-gene insertion inadvertently silenced the expression of the recombinant allele to create haploinsufficient mice (Haplo). Although Haplo heterozygotes are viable and fertile, they have ∼50% lower levels of VPS35 compared to their wild type littermates. Double heterozygous crosses fail to breed to homozygosity as a complete loss of retromer is embryonic lethal^28^ (and unpublished data). Haplo animals crossed with transgenic mice expressing Cre recombinase successfully excised the floxed mini-gene cassette to produce animals that express VPS35 p.D620N (VKI). Both strains have been maintained on the same C57Bl/6J background for >10 generations. All breeding, housing, and experimental procedures were performed according to the Institutional Animal Care and Use Committee at University of Florida. All mice were kept on a reverse cycle (light on from 8:30pm to 8:30am) and single-sex group-housed in enrichment cages after weaning at post-natal day 21. For all experiments, 3-5-month-old male animals were used. Ear notches were taken for DNA extraction and genotype validation.

### Gene and protein nomenclature

Gene nomenclature is consistent with published guidelines (https://useast.ensembl.org/index.html). Specifically, lower-case letters are given to mouse genes whereas upper case letters are used exclusively for human genes. Protein nomenclature is consistent with Uniprot. Specifically, upper case letters are used for both human and mouse protein, with the exception of Rab proteins, which appears as lower case on uniprot and many other publications.

### Perforated patch clamp electrophysiology

Mice were anesthetized with 5% isoflurane, and then transcardially perfused with ice cold cutting solution containing (in mM): 205 sucrose, 2.5 KCl, 1.25 NaH_2_PO_4_, 7.5 MgCl_2_, 0.5 CaCl_2_, 10 glucose, 25 NaHCO_3_, and 1 kynurenic acid, saturated with 95% O_2_ and 5% CO_2_ (∼305 mOsm/kg). Horizontal slices were prepared in the same cutting solution with a vibratome (Leica VT1200S) and incubated for 45min-1hr at 32°C in a solution containing (in mM): 126 NaCl, 2.5 KCl, 1.2 NaH_2_PO_4_, 1.2 MgCl_2_, 2.4 CaCl_2_, 11 glucose, and 25 NaHCO_3_, saturated with 95% O_2_ and 5% CO_2_ (pH 7.4, ∼300 mOsm/kg). Recordings were made at 33-34°C in the same solution perfused at 2 ml/min.

Perforated recordings were made using patch pipettes with an impedance of 3-7MΩ when filled with internal solution containing (in mM) 125 K-Gluconate, 4 NaCl, 10 Hepes, 4 Mg-ATP, 0.3 Na-GTP and 10 tris-phophocreatine (pH 7.3, ∼280 mOsm/kg). Amphotericin was freshly diluted in internal solution at 195 nM concentration and used as a pore-forming agent for perforated recordings. Junction potentials were not corrected for all recordings. Cells were visualized using an upright microscope (Olympus OX50WI) with infrared/differential interference contrast optics at 40x magnification. Dopaminergic neurons were identified based on their spontaneous firing (1-5 Hz) with broad action potentials (Aps) > 1.5 ms and sag potential elicited by hyperpolarizing current injection (∼-10 mV with -100 pA current injection).

### Fast scan cyclic voltammetry

Mice were sacrificed by rapid decapitation and 300 µm coronal slices containing the striatum were cut in ice-cold cutting solution (containing in mM: 130 NaCl, 10 glucose, 26 NaHCO_3_, 3 KCl, 5 MgCl_2_, 1.25 NaH_2_PO_4_, and 2 CaCl_2_), saturated with 95% O_2_ and 5% CO_2_ (∼305 mOsm/kg, pH 7.2–7.4). Subsequently, the striatal slices were incubated at room temperature for >1 hr in an artificial cerebrospinal fluid (ACSF) (containing in mM: 130 NaCl, 10 glucose, 26 NaHCO_3_, 3 KCl, 1 MgCl_2_, 1.25 NaH_2_PO_4_, 2 CaCl_2_), saturated with 95% O_2_ and 5% CO_2_ (∼300 mOsm/kg, pH 7.2–7.4). Recordings were made at 27–29°C in the same solution perfused at ∼2.5 ml/min. Slices were visualized using an upright microscope (Olympus OX50WI) with infrared/differential interference contrast optics. Single pulse stimuli (150 µs duration) were delivered by nickel-chromium bipolar electrode (made in house) placed in the dorsolateral striatum isolated (A365, World Precision Instrument) and controlled with Clampex software. Electrically evoked dopamine responses were monitored using carbon fiber electrodes (Invilog, diameter: 32 µm, sensitivity: >20 nA/µM) placed ∼100 µm from the stimulating electrode. Triangular waveforms (ramp from −400 mV to 1200 mV to −400 mV, 10 ms duration at 10 Hz) were used to detect the oxidation and redox peaks for dopamine between 700 and 800 mV and voltametric responses were recorded, standardized and analyzed with an Invilog Voltammetry system and software (Invilog Research Ltd., Finland). The input/output paradigm consisted of single pulses of increasing intensities (50–700 μA, delivered every 2 min/ 0.0083 Hz) to determine the input required to evoke ∼70% of the maximum response, which was used for the rest of the experiment. Five single pulses were delivered at 0.0083 Hz to calculate an average dopamine transient to be used for a more accurate representation of the decay characteristics. At the end of each recording session, a 3-point calibration of each carbon fiber electrode was conducted (final concentrations: 0.5, 1.0 and 2.0 μM dopamine in ACSF).

### Amphetamine treatment and open field behavior

Amphetamine (d-Amphetamine Sulfate, A5880) was made fresh each day in sterile saline at 0.5 mg/ml. Individual animals were weighed and placed into an open field chamber (19 x 19 x 19 inch) and allowed to habituate for 30 min followed by subcutaneous injection of amphetamine at 2 mg/kg. Total distance traveled was recorded throughout habituation and for 60 min following amphetamine injection. Distance traveled was analyzed by custom-made matlab code.

### MLi-2 treatment

Mice were randomly assigned for MLi-2 (Tocris) or vehicle treatment (Captisol), delivered once daily via intraperitoneal (IP) injection for a total of 7 days. Mice were weighed daily before injection. A solution of 0.5 mg/mL MLi-2 in 45% Captisol was prepared for injection, with a vehicle control of 45% Captisol alone, and filter sterilized prior to use. Each mouse was weighed and injected with a dose of 5 mg/kg. The mice were sacrificed 1 hr after the final injection, and their brains were extracted for FSCV or immunoblotting.

### Immunoblotting

Striatal tissues were homogenized and lysed in 1x Lysis Buffer (Cell Signaling #9803S) with 1 mM PMSF and 1X phosphatase inhibitors (Thermofisher; #36978 and #78427, respectively). Homogenates were spun at 12,000 g for 15 min at 4 °C and pelleted debris were removed. Samples were denatured with 5% 2-mercaptoethanol in 1x NuPage LDS sample buffer (#NP0008), and resolved by SDS-PAGE using NuPage 4–12% Bis–Tris gel (Thermo Fisher Scientific #NP0321BOX) and transferred to PVDF membranes (EMD Millipore). The blots were blocked for 1 h in 5% non-fat milk (or BSA for phosphoprotein) and incubated overnight with primary antibodies in blocking buffer at 4°C. All antibody concentrations used are detailed in Table S1. Following 3 x 15 min washes with TBS-T, the membrane was incubated at room temperature with secondary antibodies diluted in blocking solution for 1 h, washed 3 x 15 min and scanned using a Chemi-doc imaging system (Cell-Bio).

### Statistics and data reporting

All experiments, data processing and analysis were conducted in a blinded manner. All electrophysiology and fast-scan voltammetry trace were analyzed using clampfit (Molecular devices). Densitometric signal from western blots was analyzed in Image J software^29^ and normalized to either GAPDH loading control or non-phosphorylated total protein level. Data are presented throughout as mean ± SEM where “n” represents the number of slices or neurons, and “N” represents the number of animals. All data were tested for normality and statistics were performed using either nonparametric or parametric tests using Prism 8.4 (GraphPad).

## Results

### LRRK2 kinase activity is differentially affected by Vps35 haploinsufficiency and the Vps35 p.D620N mutation

We generated Vps35 haploinsufficient mice (Haplo) and Vps35 knock-in mice (VKI) that constitutively express the Vps35 p.D620N mutation, genetically linked to PD^7^ (Materials and Methods, Fig. 1A). Haplo animals exhibit a 50% reduction in Vps35 expression (Fig.1B), whereas hetero- and homozygous VKI animals exhibit comparable expression of Vps35 compared to wild type (WT) littermates (Fig. 1C)^15^. Both Haplo and VKI animals are viable and fertile, whereas homozygous Vps35 knockout is embryonic lethal^28^. Haplo animals are a *bona-fide* loss of Vps35 function model. Thus, we performed a comparative examination of VKI and Haplo animals at 3-months of age to investigate gain-versus loss-of-function of VPS35 with respect to LRRK2 biology, and assess the interplay of VPS35-LRRK2 in the dopamine system.

**Fig. 1:**
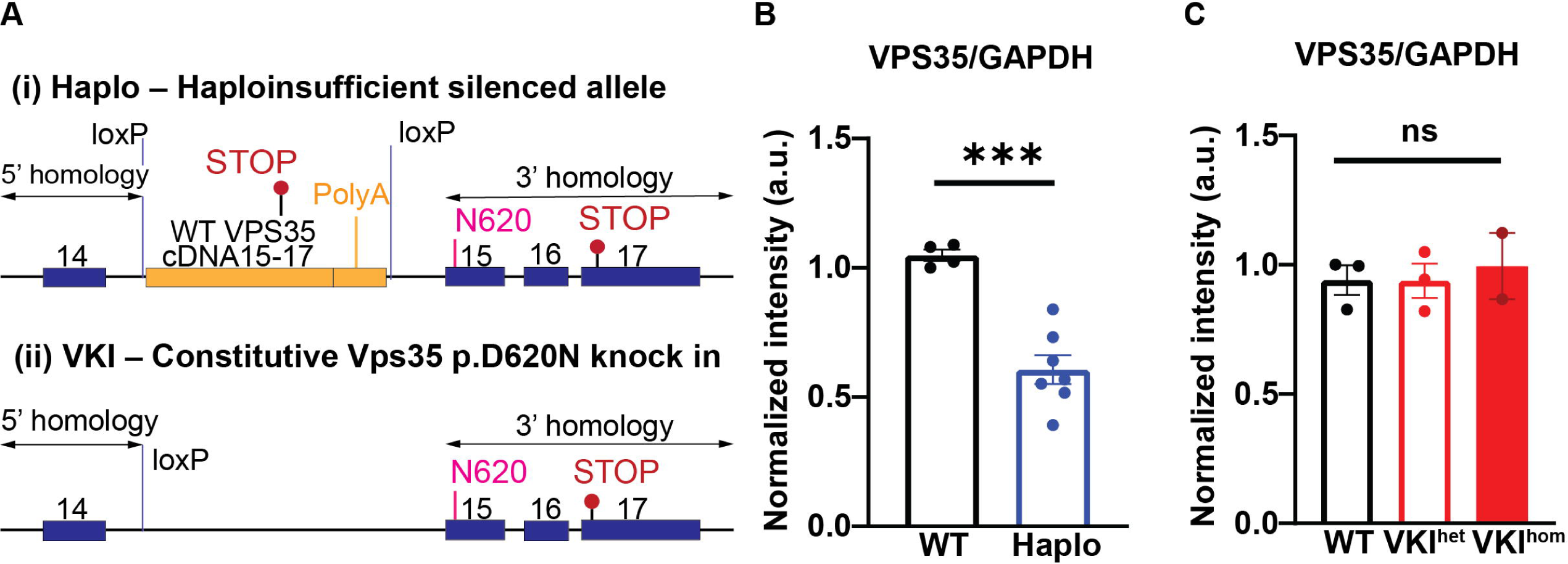
Generation of Vps35 haploinsufficient (Haplo) and Vps35 p.D620N knock-in (VKI) mice. **A:** Schematics of targeting design for **(i)** Haplo animals showing the murine Vps35 genomic sequence (Ensembl reference ENSMUSG00000031696), 5′ and 3′ homology arms (arrowed), exons 14–17 (blue), engineered loxP site in intron 14, the g.85,263,520G>A mutation in exon 15 (encoding the p.N620 substitution, in pink) and endogenous stop codon in exon 17 (TAA, lollipop in red). A cDNA cassette of wild type murine ‘exons 15-17 with a polyA tail’ (yellow) was originally introduced to create conditional knock-in (VKI), but inadvertently silenced the endogenous allele. **(ii)** VKI animals created by Cre-recombinase deletion between loxP sites. **B-C:** Total VPS35 protein level were quantified by western blotting, showing 50% reduction in haplo compared to their WT littermates. (**B**: WT: N=4, Haplo: N=7; unpaired t test, t_9_=5.760, p<0.001) and no difference in Het and Hom VKI (**C**: WT: N=3, Het: N=3, Hom: N=2; One-way ANOVA, F_2, 5_= 0.15, p=0.87). *p < 0.05, **p < 0.01, ***p < 0.001. Error bars are ±SEM.

We first performed a cursory examination of the endo-lysosomal pathway by western blot analysis of striatal homogenates from VKI and Haplo mice. We used Rab10 (pThr73), Rab12 (pSer106) and Rab29 (pThr71) as indirect readouts of LRRK2 kinase activity and as markers for endosomal, lysosomal and Golgi trafficking, respectively^23, 30–32^. We found Haplo animals exhibit significant reduction in pRab12/Rab12 (Fig. 2A, D) and comparable levels of pRab10/Rab10 (Fig. 2C) and pRab29/Rab29 (Fig. 2E) compared to WT littermates. Previous work has shown LRRK2 kinase activation in VKI^23^. Here, we confirmed an increase in pRab10 (Fig. 2F) and pRab12 (Fig. 2G) and showed that pRab29/Rab29 is also significantly increased in striatal homogenates from 3-month-old VKIs (Fig. 2H). Treatment with MLi-2 *in vivo* (7-day intraperitoneal injection, 5mg/kg dose), a potent LRRK2 kinase inhibitor, significantly abolished Rab phosphorylation in VKIs with a clear interaction between treatment and genotype, further supporting that hyperphosphorylation of Rab proteins in VKIs are LRRK2 kinase dependent (Figure S1). Rab12 is recently identified to recruit and activate LRRK2 to damaged lysosomal membrane. Upon examining down stream lysosomal and autophagic markers, we observed a significant increase in lysosomal associated protein-1 (LAMP1) in both models (Fig. 1F, J). While both Vps35 haploinsufficiency and the p.D620N mutation lead to endo-lysosomal deficits, the data suggests p.D620N induces a gain-of-function with respect to LRRK2 kinase activity.

**Fig. 2:**
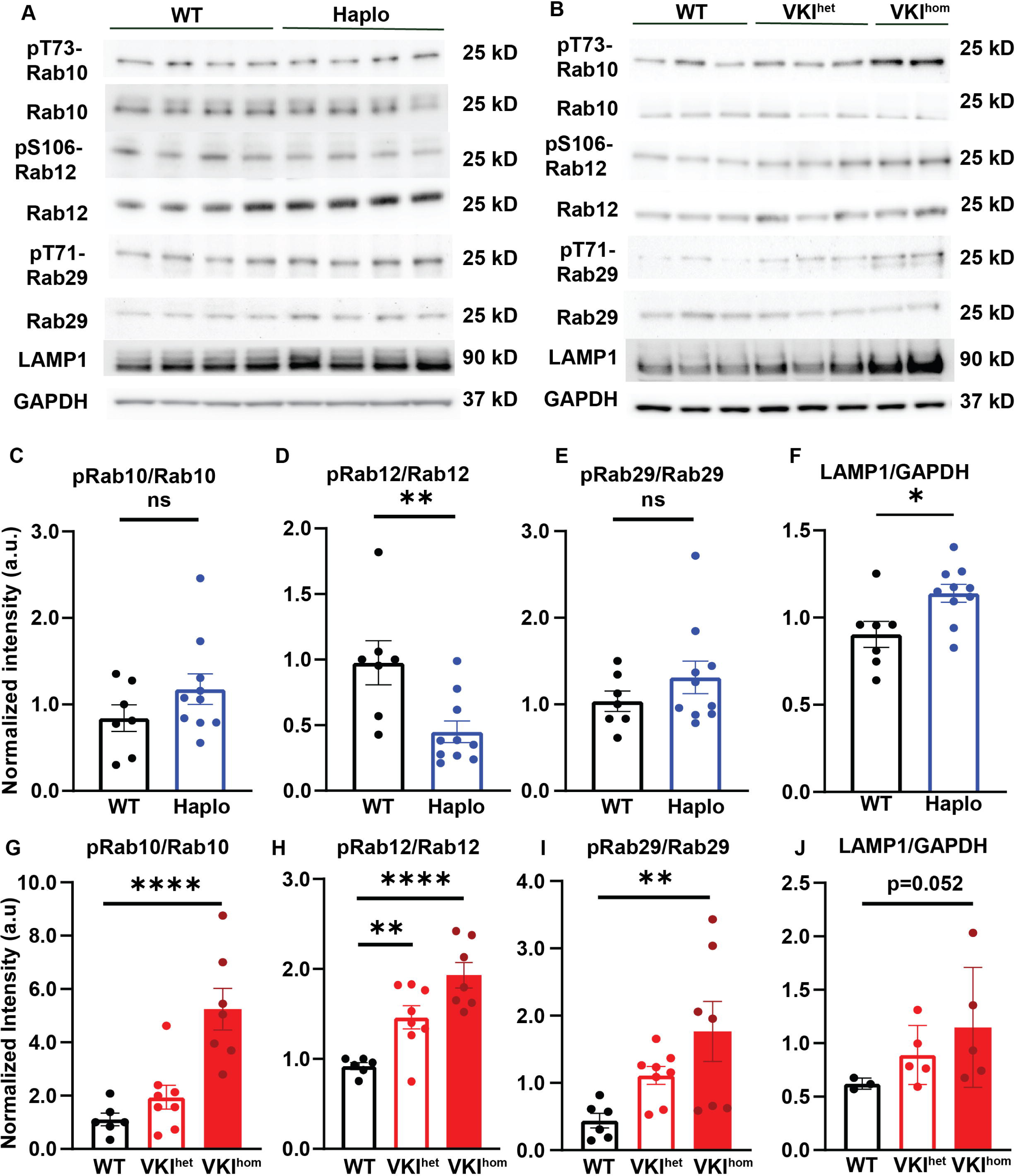
Rab protein phosphorylation is reduced in Haplo but increased in VKI. **A-B:** Representative blots of Rab protein phosphorylation in striatal lysate of Haplo and VKI animals respectively. **C-E**: Quantifications of phospho/ total protein levels show Haplo animals exhibit significant reduction in phospho-Rab12 (**D**: WT: N=7, Haplo N=10, unpaired t-test; t_15_ =3.09, p<0.001) and comparable level of phospho-Rab10 (**C**: WT: N=7, Haplo N=10, unpaired t-test; t_15_ =1.35, p=0.19) and phospho-Rab29 (**E**; WT: N=7, Haplo N=10, unpaired t-test; t_15_ =1.12, p=0.28). **F**: Haplo animals also exhibit significant increase in LAMP1 expression (unpaired t test, t_15_=3.17, p<0.01) **G-J**: VKI exhibit significant increase in phospho-Rab10 (**G:** WT: N=6, VKI^het^: N=8, VKI^hom^: N=7; One-way ANOVA F_2, 18_ =15.14, p=0.0001; Dunnett’s multiple comparison test; WT vs. VKI^het^ : p=0.48; WT vs. VKI^hom^ : p=0.0002), phospho-Rab12 (**H**: WT: N=6, VKI^het^ : N=8, VKI^hom^ : N=7; WT vs. Het: p=0.01; One-way ANOVA: F_2, 18_ =16.42, p<0.0001; Dunnett’s multiple comparison test; WT vs. VKI^hom^ : p<0.0001), phospho-Rab29 (**I**: WT: N=6, VKI^het^ : N=8, Hom: N=7; One-way ANOVA; F_2, 18_ =5.22, p<0.05; Dunnett’s multiple comparison test; WT vs. Het: p=0.18; WT vs. Hom: p<0.01) and LAMP1 (**E**: WT: N=6 animals; VKI^het^: N=8 animals; VKI^hom^: N=7 animals; One-way ANOVA: F_2, 18_=3.20, p=0.06; Dunnett’s multiple comparison test: WT vs. VKI^hom^ p=0.05) *p < 0.05, **p < 0.01, ***p < 0.001. Error bars are ±SEM.

### Vps35 haploinsufficiency or p.D620N mutant dysfunction has minimal effect on basal dopaminergic neuron pace-making activity

The loss of dopaminergic neurons in the *substantia nigra pars compacta* (SNpc) region leads to the cardinal motor symptoms of PD. To examine how dysregulation of VPS35 affects SNpc dopaminergic neurotransmission, we used *ex-vivo* perforated patch clamp electrophysiology to examine autonomous pace-making activity, an energy-demanding process that may contribute to the selective vulnerability of these neurons^33, 34^. SNpc dopaminergic neurons are identified based on their large cell body, their spontaneous firing (1-5 Hz), broad action potentials (APs) > 1.5 ms (Fig. 3A), and prominent depolarizing sag during hyperpolarizing current injection (Fig. 3B)^35^. The sag potential in SNpc dopaminergic is regulated by hyperpolarization-activated cyclic nucleotide-gated (HCN, I_h_) cation channels, an important regulator in pace-making activity^35, 36^. Both Haplo and VKI^het^ dopamine neurons exhibit comparable levels of sag potential, reflecting normal HCN function (Fig. 3C). The distribution of interspike intervals (ISI) is commonly used to study the firing pattern. Although Haplo neurons exhibit a significant right-ward shift of ISI at the level of each action potential (Fig. 3D), the mean firing frequency at cellular level (Fig.3E) and the fidelity (Fig. 3F) of pace-making activity is comparable among dopaminergic neurons of WT, Haplo and VKI^het^. Together, these results suggest that the pace-making activity of SNpc dopaminergic neurons is intact in 3-month-old Haplo and VKI^het^ animals.

**Fig. 3:**
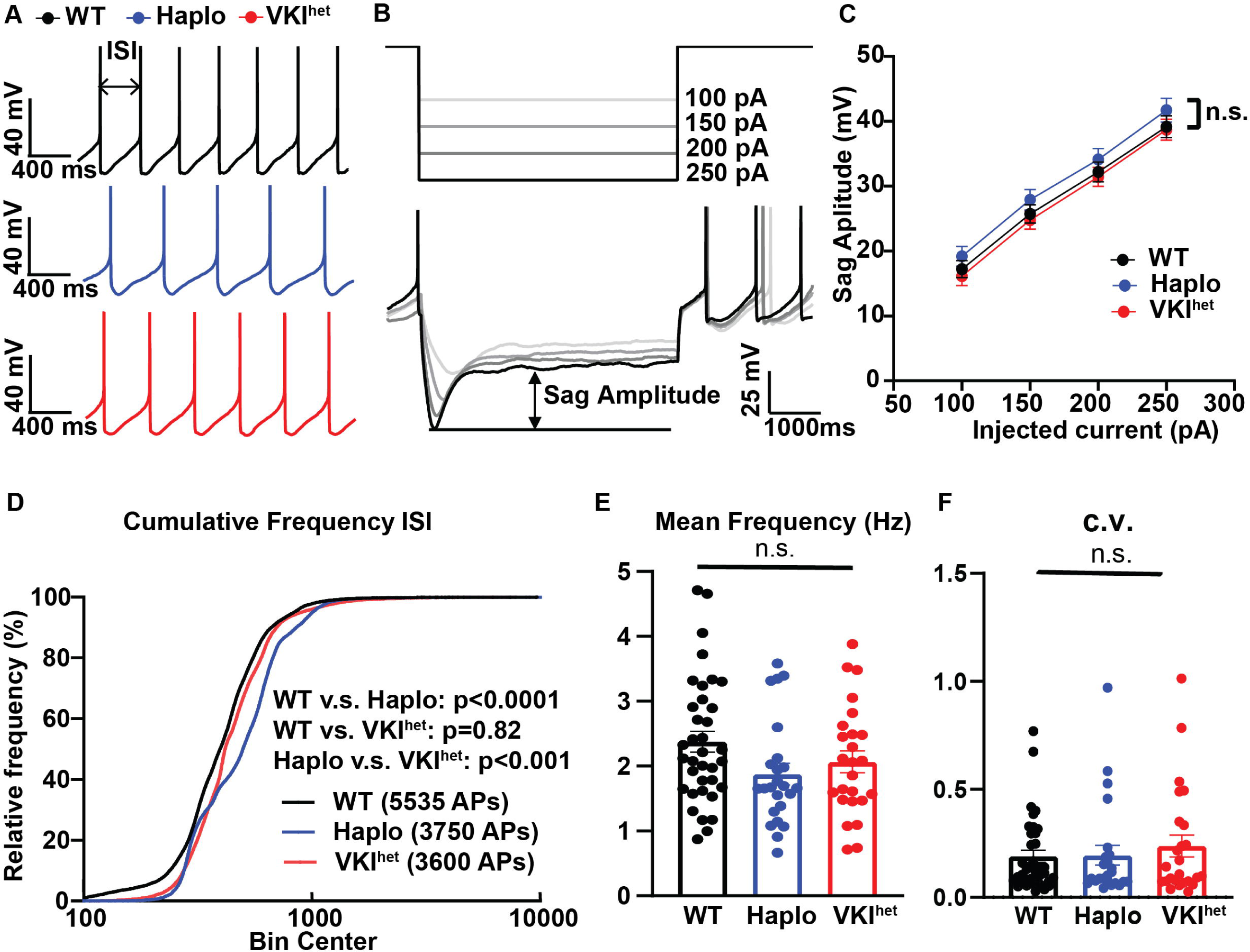
Haplo and VKI SNc dopaminergic neurons exhibit comparable basal pace-making activity. **A:** Representative traces of perforated, spontaneously active SNpc dopaminergic neurons. **B:** Representative traces displaying depolarizing sag potentials in response to hyperpolarizing current injections in SNpc dopaminergic neurons. **C:** Summary plot of sag amplitude show comparable I_H_ activation in VKI^het^ and Haplo (Two-way ANOVA; WT: n=34, N=7, VKI^het^ n=25, N=5, Haplo: n=24, N=5; Genotype: p>0.05). **D**: Cumulative frequency plot of inter-spike intervals (ISIs) of all the action potentials (APs) shows significant increased ISI in Haplo compared to WT and VKI^het^. (WT: n=23,822 APs, Haplo: n=13,492 APs, VKI^het^: n=15,213 APs, One-way ANOVA with Tukey’s multiple comparisons test; WT vs. Haplo: p<0.0001; WT vs. VKI^het^; p=0.82; Haplo vs. VKI^het^: p<0.001). **E-F**: VKI^het^ or Haplo exhibit no significant difference in firing frequency (**E**: WT: n=34 cells, N=7 animals, VKI n=25 cells, N=5 animals, Haplo: n=24 cells, N=5 animals; One-way ANOVA, F_2, 82_ =2.37, p=0.10) or the fidelity of the pace-making activity as reflected by coefficient of variance (C.V.) of all ISIs. (**F**: WT: n=34 cells, N=7 animals, VKI n=25 cells, N=5 animals, Haplo: n=24 cells, N=5 animals, Kruskal-Wallis test, p=0.82). *p < 0.05, **p < 0.01, ***p < 0.001. Error bars are ±SEM.

### Haplo and VKI animals exhibit similar dysfunction in striatal dopamine release

SNpc dopaminergic neurons project extensively to the dorsal lateral striatum. Dopamine release at striatal axonal terminals critically regulates sensorimotor behavior and activity, independent of the pace-making of soma in the midbrain^37–39^. To examine the effects of VPS35 dysfunction in striatal dopamine dynamics, we performed *ex-vivo* fast scan cyclic voltammetry in acute striatal slices of Haplo and VKI animals. Upon electric stimulation, Haplo slices exhibit elevated peak amplitude in dopamine release across a broad range of stimulation intensity (Fig. 4A-B), and a significant increase in the decay constant tau (Fig. 4C), reflecting prolonged dopamine reuptake. This response phenocopied age-matched striatal slices of VKI, which also exhibit enhanced peak amplitude and prolonged reuptake kinetics (Fig. 4D-F). Dopamine reuptake in the dorsal striatum is primarily mediated by the dopamine transporter (DAT), and both Haplo and VKI mice exhibited a significant reduction in striatal DAT expression, which is consistent with prolonged dopamine reuptake kinetics in both strains (Fig 4.G-I). Collectively, these results are consistent with previous literature showing DAT as a retromer cargo^40^, and provide complementary *in-vivo* evidence that dysregulation of retromer impairs DAT trafficking, which in turn leads to dysregulation of striatal dopamine dynamics.

**Fig. 4:**
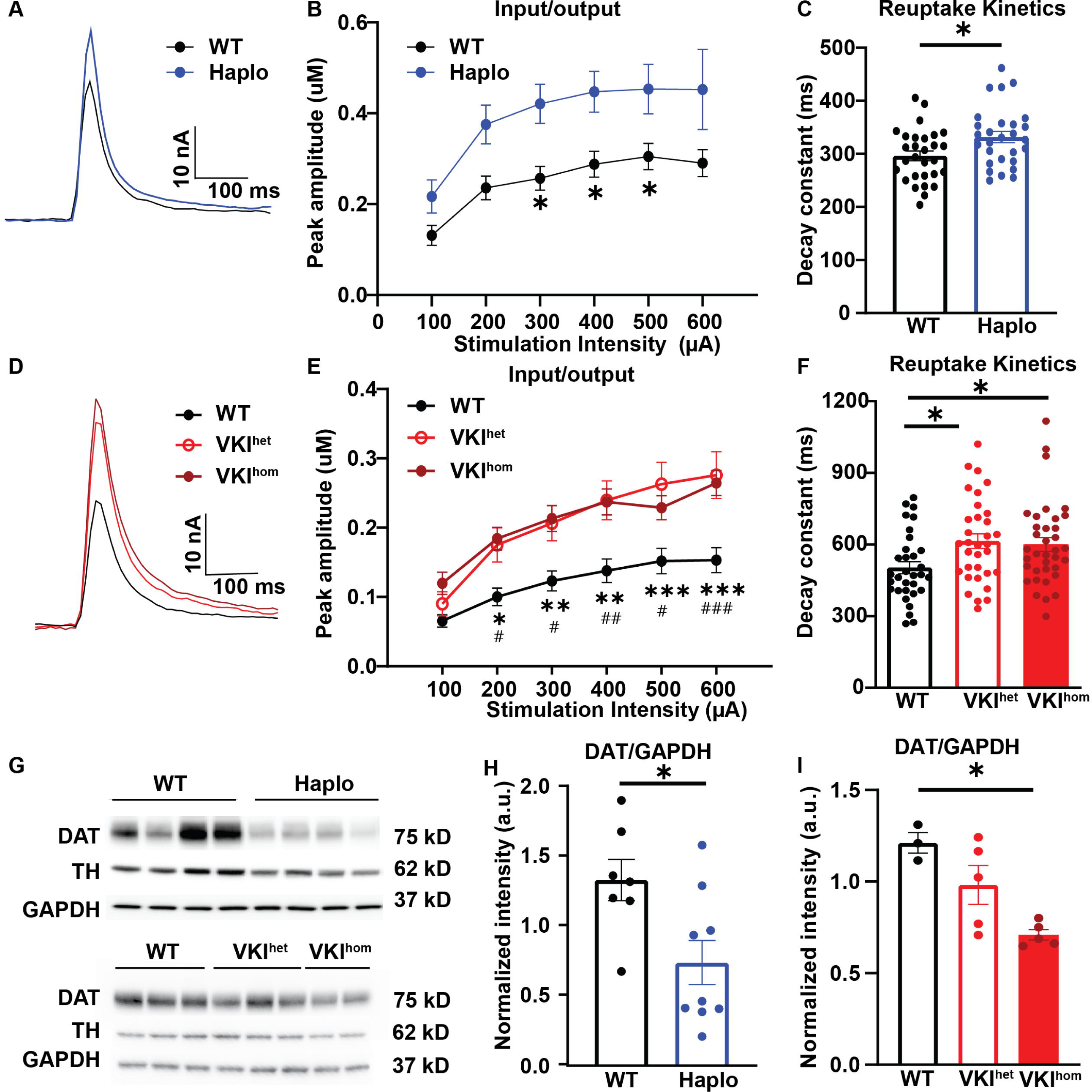
Haplo and VKI mice exhibit increased evoked peak amplitude, reduced DA reuptake and reduced DAT expression. **A, D**: Representative traces of evoked dopamine response following single pulse stimulation in Haplo and VKI animals respectively. **B-C:** Haplo slices exhibit significant increase in evoked peak amplitude as reflected by Input/output curves (**B**: WT n=29 slices. N=6; Haplo n=28 slices, N=5 animals; Mixed effects ANOVA with Sidak’s multiple comparisons test, stimulation intensity: F_(5, 253)_ = 28.33, p<0.0001; genotype: F_(1, 55)_ = 8.20, p<0.01; stimulation intensity x genotype F_(5, 253)_ = 1.02, p=0.41. WT v.s. Haplo: *p < 0.05, **p < 0.01, ***p < 0.001) and prolonged dopamine reuptake kinetics as reflected by increased decay constant *τ* (**C:** WT n=29 slices. N=6; Haplo n=28 slices, N=5 animals; unpaired t test, t_55_=2.53, p<0.05). **E-F**: VKI exhibit significant increase in evoked peak amplitude as reflected by Input/output curve (**E:** Mixed effects ANOVA with Sidak’s multiple comparisons test, stimulation intensity: F_6, 588_ = 87.54, p < 0.0001; genotype: F_2, 98_ = 7.097, p < 0.05; stimulation intensity x genotype F_12, 588_ 3.87, p < 0.0001; Bonferroni post-test WT vs VKI^het^ *p < 0.05, **p < 0.01, ***p < 0.001, WT vs VKI^hom^; # p < 0.05, ## p < 0.01, ### p < 0.001) and prolonged dopamine reuptake kinetics as reflected by increased decay constant *τ* (**F**: One-way ANOVA F_2,98_ = 4.37, p < 0.02, Dunnett’s multiple comparison test, t_98_ = 2.71, p < 0.05, and t_98_ = 2.42, p < 0.05 for VKI^het^ and VKI^het^ slices). (**D-F** adapted from Caltaldi et al., 2018) **G:** Representative western blots of dopaminergic markers in striatal tissue. **H-I:** Quantification of western blot shows significant reduction in total striatal DAT expression in Haplo (**H**, WT: N=7, Haplo: N=10; unpaired t test; t_15_=2.62, p<0.05). and VKI mice (**I**, WT N=3, VKI^het^: N=5, VKI^hom^: N=5; One-way ANOVA, F_2, 10_ =9.53, p<0.01; Dunnett’s multiple comparisons test; WT vs. VKI^het^ p=0.13, WT vs. VKI^hom^, p< 0.01). Data normalized to GAPDH loading control. *p < 0.05, **p < 0.01, ***p < 0.001. Error bars are ±SEM.

### Haplo and VKI animals exhibit divergent locomotion responses to amphetamine

Nigro-striatal dopamine neurotransmission critically regulates locomotion. We and others have previously characterized motor function in Haplo and VKI mice at 3 months of age, but found no significant differences in open field exploration, rotarod or cylinder test at baseline^13, 15, 28^. Given the constitutive nature of our genetic model, it is possible that the neuronal network has adapted to subtle changes in dopamine transmission, which could mask behavioral responses. Therefore, we acutely challenged the dopamine system with amphetamine (AMPH), a potent stimulator of dopamine release via DAT reverse transport and vesicular depletion, and measured AMPH induced locomotion in Haplo and VKI^het^ mice. Animals were placed in an open field chamber for 30 min prior to subcutaneous injection of AMPH at 2 mg/kg, and then monitored for an additional 60 min within the open field chamber. AMPH induced hyperlocomotion in WT, VKI^het^ and Haplo; however, Haplo mice exhibited a reduced response compared to WT and VKI^het^ mice (Fig. 5 A-B), consistent with the effects of AMPH treatment in DAT knockout mice^41, 42^. In contrast, although VKI^het^ mice also have reduced DAT reuptake, they exhibit a rapid and robust increase in their locomotion, and initially exceeding that observed in WT. Hence, Vps35 p.D620N confers a gain-of-function to maintain AMPH-induced locomotion.

**Fig. 5:**
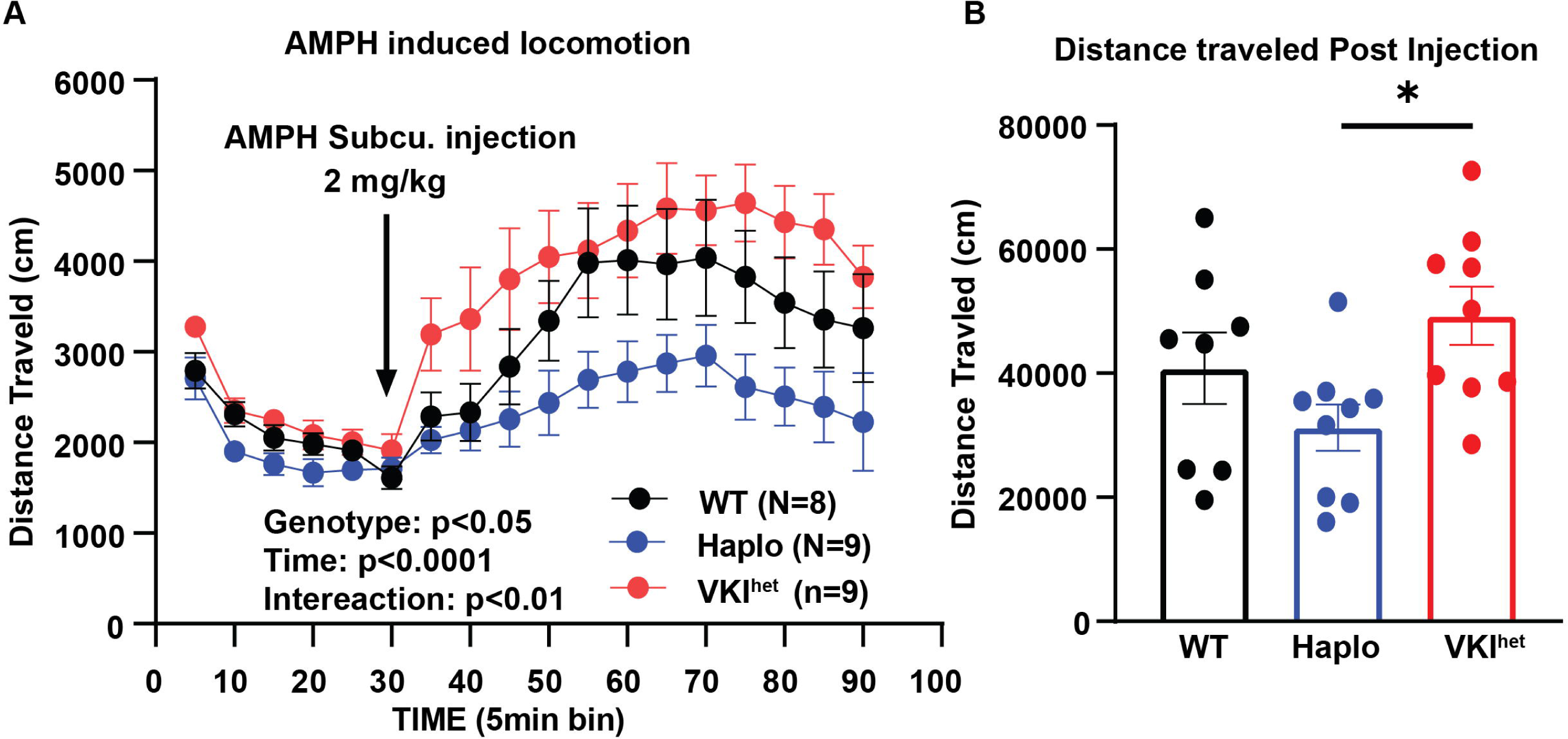
Haplo and VKI exhibit divergent responses to amphetamine induced locomotion. **A**: Distance traveled over 90 min of open field monitoring. Haplo and VKI^het^ animals exhibit divergent responses to AMPH (WT: N=8 animals, Haplo N=9 animals, VKI^het^ N=9 animals; Two-way ANOVA: Genotype: F_2, 23_ =5.04, p<0.05; Time: F_2.483, 57.11_ =22.41, P<0.0001; Time x Genotype F_34, 391_ = 1.76, p<0.01; Subject: F_23, 391_ = 20.78, p<0.0001) **B**: Total distance traveled post injection. (WT: N=8 animals, Haplo N=9 animals, VKI^het^: N=9 animals; One-way ANOVA; F_2, 22_=5.14, p<0.05; Tukey’s multiple comparison test Haplo vs. VKI^het^: p<0.05). *p < 0.05, **p < 0.01, ***p < 0.001. Error bars are ±SEM.

### Pharmacological inhibition of Lrrk2 kinase activity *in-vivo* rescues dopaminergic phenotypes in VKI

Next, we investigated whether the divergent responses in AMPH-induced hyperlocomotion in Haplo and VKI^het^ are dependent on LRRK2 activity. We acutely inhibited LRRK2 kinase activity *in-vivo* using MLi-2, a potent LRRK2 kinase inhibitor. 3-month-old animals received either vehicle or MLi-2 treatment for 7 days through intraperitoneal injection (Fig. 6A, Fig. S2). Interestingly, while MLi-2 treatment has minimal effects on WT and Haplo mice, it attenuated AMPH-induced hyperlocomotion in VKI^het^ animals to levels comparable to Haplo (Fig. 6), suggesting that AMPH-induced hyperlocomotion in VKI^het^ is dependent on LRRK2 kinase activation.

**Fig 6:**
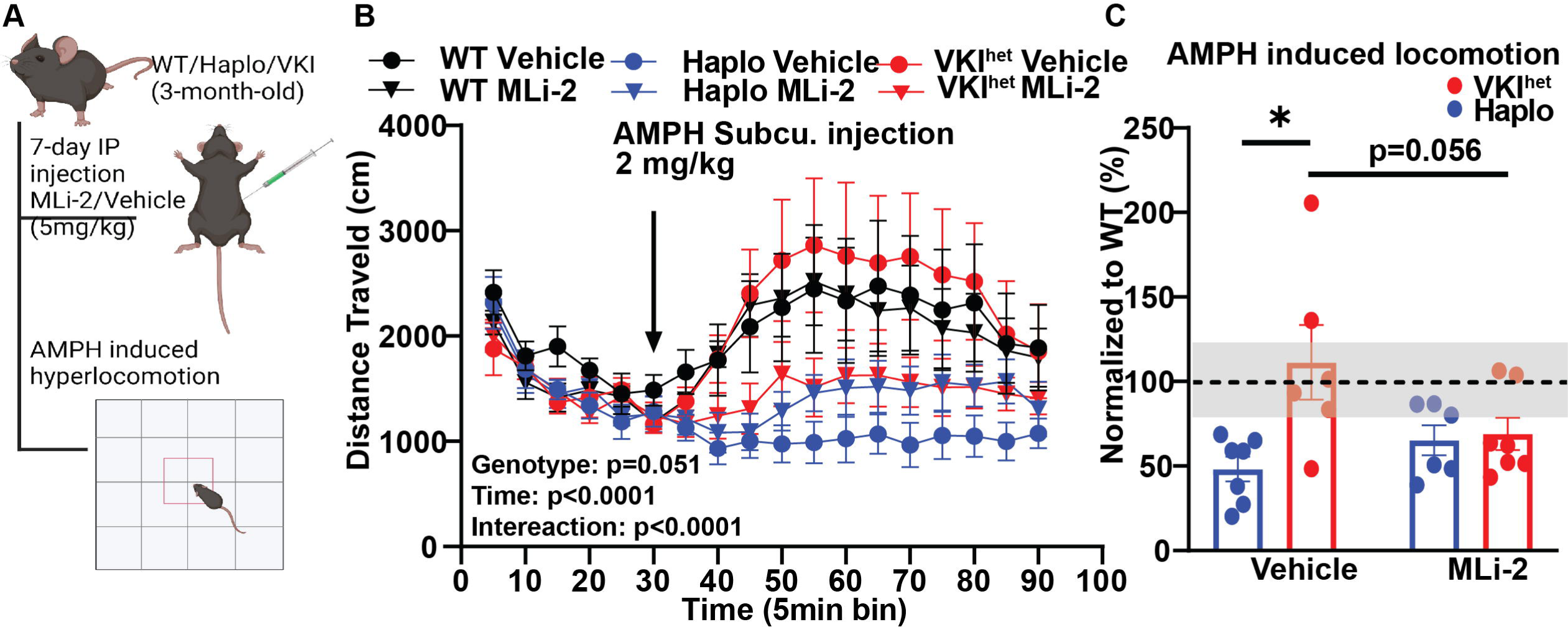
7-day *in-vivo* treatment by MLi-2 reduces AMPH induced hyperlocomotion in VKI to levels comparable to Haplo mice. **A:** Schematic of MLi-2 treatment and experimental paradigm (Created with BioRender). **B:** Distance traveled over 90 min of open field monitoring (WT vehicle, N=7, WT MLi-2: N=7, Haplo Vehicle: N=7, Haplo MLi-2: N=6, VKI^het^ vehicle: N=6, VKI^het^ MLi-2: N=7; Two-way ANOVA: Group: F _5, 33_ = 2.48, p=0.051; Time: F _2.053, 67.76_ =9.33, p<0.001; Time x Group: F _85, 561_ =2.115, p<0.0001; Subject F_33, 561_ =22.78, p<0.0001). **C:** Total distance traveled post injection. Initial elevation in VKI^het^ was abolished by MLi-2 treatment. Data are normalized by average of WT Vehicle group (Dotted line, gray bar represent WT average ±SEM) (WT veh, N=7, WT MLi2: N=7, Haplo Vehicle: N=7, Haplo MLi-2: N=6, VKI^het^ vehicle: N=6, VKI^het^ MLi-2: N=7; Two-way ANOVA: Genotype: F _(1, 22)_=0.98, p=0.33; Treatment F _1, 22_ = 6.88, p<0.05; Genotype x Treatment F_1, 22_ = 5.40, p<0.05; Sidak multiple comparisons: VKI^het^ vehicle v.s. VKI^het^ MLi-2 p=0.056) *p < 0.05, **p < 0.01, ***p < 0.001. Error bars are ±SEM.

Phosphorylation locks Rab proteins in their GTP-bound forms, prevents their interaction with effector proteins and stalls vesicular trafficking. To test whether LRRK2 mediated hyperphosphorylation of Rab proteins impairs DAT trafficking in VKI^het^, we re-examined evoked striatal dopamine dynamics in VKI^het^ mice treated with vehicle or MLi-2 (Fig. 7A).

**Fig. 7:**
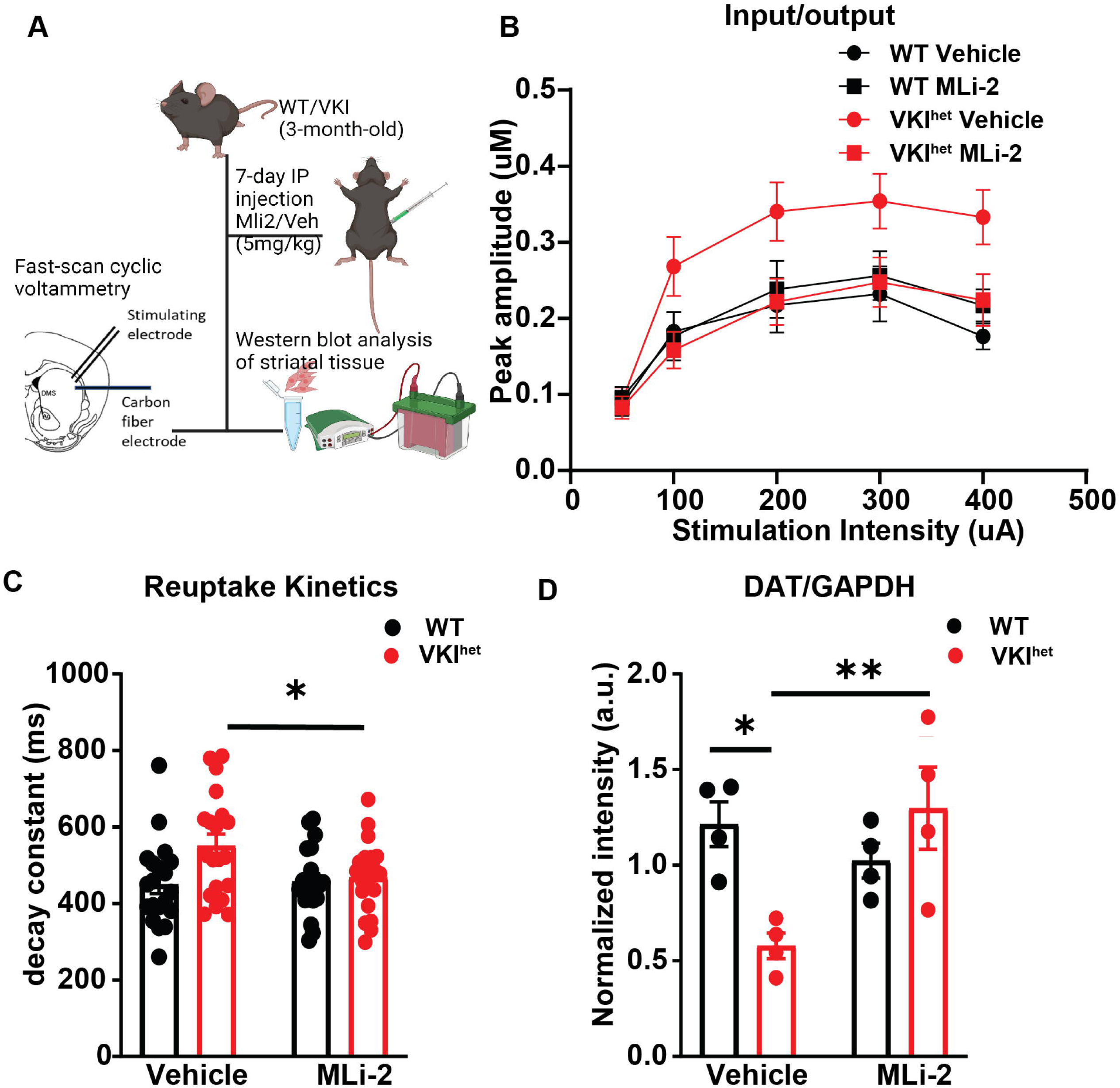
7-day *in-vivo* treatment by MLi-2 normalizes evoked striatal dopamine response and Dat expression in VKI. **A:** Schematic of MLi-2 treatment and experimental paradigm (Created with BioRender). **B:** MLi-2 treatment normalizes input/output response of VKI^het^ to levels comparable to WT (WT vehicle: n=19 slices, N=5 animals, WT MLi-2: n=20 slices, N=5 animals, VKI^het^ vehicle: n=20 slices, N=5 animals, VKI^het^ MLi-2: n=20 slices, N=5 animals; Mixed effect analysis: stimulation intensity: F_4, 184_=21.99, p<0.0001; Genotype: F_1, 184_=9.89, p<0.01; Treatment: F_1, 161_ =7.57, p<0.01; Genotype x Treatment: F_1, 161_ =16.57, p<0.0001) **C:** MLi-2 treatment normalizes reuptake kinetics of VKI^het^ to level comparable to WT. (WT vehicle: n=19 slices, N=5 animals, WT MLi-2: n=20 slices, N=5 animals, VKI^het^ vehicle: n=20 slices, N=5 animals, VKI^het^ MLi-2: n=20 slices, N=5 animals; Two-Way ANOVA: Genotype, F_1, 76_=5.33, p<0.05; Treatment, F_1, 76_ =2.40, p=0.13; Genotype x Treatment: F_1, 76_=3.49, p=0.066; Tukey’s multiple comparisons: VKI^het^ Vehicle vs VKI^het^ MLi-2: p<0.05). **E:** MLi-2 treatment normalizes DAT expression level in VKI^het^. (N=4 for all groups; Two-Way ANOVA: Genotype, F_1, 12_ =1.81, p=0.20; Treatment: F_1, 12_=3.84, p=0.074; Genotype x Treatment: F _1, 12_ =11.39, p<0.01; Tukey’s multiple comparisons: VKI^het^ Vehicle vs VKI^het^ MLi-2: p<0.05; VKI^het^ Vehicle vs. WT Vehicle: p<0.05) *p < 0.05, **p < 0.01, ***p < 0.001. Error bars are ±SEM.

Remarkably, MLi-2 treatment normalized dopamine dynamics in VKI^het^ to levels comparable to WT, both in terms of peak amplitude (Fig. 7B) and reuptake kinetics (Fig. 7C). At the protein level, the initial reduction in striatal DAT expression in VKI^het^ animals was also raised to levels comparable to WT (Fig. 7D). Taken together, these results suggest that LRRK2 functions downstream of VPS35 to regulate DAT trafficking and dopamine signaling.

## Discussion

Deficits in endo-lysosomal trafficking are shown to link multiple PD-related genes and may disproportionally affect dopaminergic neurons due to their extensively arborized axons and requirements for nigrostriatal axonal trafficking^43, 44^. Here we show that Vps35 p.D620N confers a gain of function with respect to LRRK2 kinase activity, and a reduction of DAT function and expression. Moreover, acute inhibition of LRRK2 kinase activity is sufficient to rescue the deficits in nigrostriatal dopamine dynamics and related AMPH-induced behavior. A strength of our work is that we are able to directly compare Haplo and VKI results, as these mouse models were derived from the same founder and have been maintained on the same genetic background (Fig. 1). By focusing on DAT, a known retromer cargo specific to dopaminergic terminals, we were able to assess the interplay between VPS35 and LRRK2 in a cell specific manner *in vivo*. Our approach was designed to be etiologically faithful and physiologically relevant to dopamine biology, which is clearly compromised in PD. Hence, results in the VKI^het^ model and potentially human subjects with VPS35 p.620N may inform the development of LRRK2 kinase inhibitors as a disease-modifying therapy.

VPS35 and LRRK2 functionally interact to regulate multiple vesicular trafficking pathways including endocytic trafficking^45, 46^, synaptic vesicle recycling^25^ and trans-Golgi network (TGN) trafficking^47^. By western blotting, we identified significant increase in phosphorylation of Rab proteins, notably pRab10, pRab12 and pRab29 in VKI compared to WT, suggesting Rab-mediated endosomal, lysosomal and Golgi trafficking may be compromised. Owing to the difficulty in detecting pS1292 LRRK2 in murine brain under basal conditions, we were not able to directly observe a significant increase in this epitope. Nevertheless, application of the LRRK2 kinase inhibitor MLi-2 significantly reduced pRabs in VKI mice with a clear interaction between treatment and genotype, suggesting Rab hyper-phosphorylation in VKI mice is LRRK2 kinase mediated. (Fig. 2). In contrast, there is little evidence for dysregulation of LRRK2 kinase activity in Vps35 haploinsufficiency, since we observed a significant reduction in pRab12/Rab12, and no changes in pRab10/Rab10 or pRab29/Rab29 (Fig. 2). The reduction in pRab12/Rab12 may reflect a compensatory increase in endo-lysosomal flux as retromer cargos are inadvertently retained, rather than efficiently recycled^48–50^. Hence, in agreement with others, we provide *in-vivo* evidence that Vps35 p.D620N results in a gain-of-function with respect to LRRK2 kinase activity^23^. Nevertheless, protein phosphorylation is also temporally labile and is more challenging to quantify from dissected tissues than cell lysates, and both these phenomena may lead to disparities in the literature. For example, we note that siRNA knockdown of Vps35 in mouse embryonic fibroblasts induces a significant reduction in pRab10/total Rab10 signal^23^.

At 3-months of age, we observed no significant change to the pace-making activity of the dopaminergic soma in the SNpc (Fig. 3) but evoked striatal dopamine release is significantly enhanced in Haplo and VKI slices, along with a significant reduction in DAT expression and function. (Fig. 4). Dopaminergic terminals in the striatum are under the control of local micro-circuitry^39, 51^ and striatal dopamine release can be decoupled from the firing activity of the soma^37, 38^. Vps35 is enriched at dopaminergic release sites, ^52^ thus it is not surprising to see that Vps35 dysfunction can lead to dysregulated dopaminergic neurotransmission in a region-specific manner. We found both Vps35 haploinsufficiency and the p.D620N mutation impairs DAT function and expression. This complements previous studies that show DAT is a cargo protein for retromer-dependent trafficking^53, 54^. Behavioral response of Haplo mice to AMPH was blunted, similar to hetero- and homozygous DAT knockout mice^41, 42^. Despite deficits in DAT, VKI^het^ exhibit enhanced AMPH induced locomotion compared to Haplo, which is abolished by MLi-2 treatment (Fig 5-6). Hence, Vps35 p.D620N confers a gain of function to maintain AMPH locomotion that is LRRK2 dependent.

DAT critically regulates dopamine neurotransmission and its function is dynamically regulated by multiple endocytic trafficking pathways^55^. A reduction in DAT density and increased dopamine turnover are observed in early symptomatic and pre-symptomatic individuals with monogenic mutations causal for parkinsonism^56–58^. Thus, early alterations in striatal dopamine dynamics and DAT expression in Haplo and VKI mice are consistent with human imaging results. Although there is no overt neurodegeneration at 3-month of age^15^, VKI mice do exhibit an age-dependent loss of midbrain dopaminergic neurons^17^, a cardinal feature of PD. Collectively, our results and prior studies highlight VKI mice as a biologically and clinically relevant model to study molecular and circuit mechanisms underlying the dysfunction of dopamine neurons before the onset of neurodegeneration.

Intriguingly, acute inhibition of LRRK2 activity also rescued striatal DAT expression and dopamine dynamics in *ex-vivo* slice preparations (Fig. 7). It’s unlikely the results of our 7-day treatment paradigm are dependent on the synthesis of new DAT. *In-situ* hybridization of DAT mRNA shows an intense signal in the midbrain and is minimal in striatum (https://mouse.brain-map.org/gene/show/12942)^59^, suggesting DAT transcriptional and translational production in the striatum is negligible. Rather, DAT synthesis occurs in the soma; subsequent anterograde axonal transport is mainly through membrane diffusion, with only a minor contribution from vesicular trafficking, and it takes around 20 days for new DAT to replenish levels in striatal dopaminergic terminals^53^. Thus, the increase in DAT in VKI mice with LRRK2 kinase inhibition more likely reflects less endosomal retention and/or autophagy in dopaminergic terminal and axons and, in support, neuronal LRRK2 kinase activity has recently been demonstrated to disrupt axonal autophagosome transport and impair acidification^60^. Prior *in vitro* studies have also shown Lrrk2 p.G2019S expression retains epidermal growth factor receptor (EGFR) in the endocytic compartment and impairs both its degradation and recycling^61^, a deficit was rescued upon expression of phosphodeficient RAB8A variant^61^. Therefore, we postulate LRRK2 kinase inhibition rescues DAT expression by facilitating the endosomal retrieval and recycling of DAT in a Rab-dependent manner^62–64^. In addition to Rab proteins, there are also reports of an interplay between LRRK2 and RIT2, a neural GTPase implicated in DAT trafficking^65, 66^. Due to technical limitations, we were unable to track intracellular DAT at an organelle level *in vivo*. Future studies using super-resolution microscopy to study DAT trafficking in dopaminergic terminals are warranted.

Our study demonstrates how VPS35 and LRRK2 orchestrate DAT localization and function, *in-vivo*, in a cargo and cell specific manner. In VKI mice the application of MLi-2 rescued DAT expression and function, and reduced the AMPH response. However, increased DAT should enhance rather than reduce AMPH sensitivity^67^. Hence, we surmise the effects of MLi-2 on AMPH induced locomotion are not mediated by changes in DAT expression and function alone. VKI mice may exhibit dysregulation in downstream circuits that are also dependent on LRRK2, which is more abundant in glia^68^, medium spiny neurons and innervating cortical synapses^69, 70^ than dopaminergic terminals or axons^69–71^. Indeed, many consider the function of LRRK2 to be as, if not more, important in immune cells including microglia. Future studies might explore this relationship in different cellular contexts by crossing VKI mice onto a conditional *Lrrk2* knockout background. Nevertheless, LRRK2 is a large multi-domain protein whose function is not limited to its kinase activity. Thus, it would be prudent to assess the role of LRRK2 scaffolding using the kinase dead p.D1994A mutant mouse. It would also be useful to compare *LRRK2* antisense oligonucleotide treatments on dopamine physiology *in vivo*, before deploying Lrrk2 kinase inhibitors as a long-term disease-modifying therapy for PD. Based on the results, dopaminergic imaging including ^18^F-DOPA turnover^71^ in human *VPS35* p.D620N heterozygotes may inform target engagement and optimal dosing for LRRK2 kinase inhibitors.

## Supplementary Figure legend

**Figure S1 (Related to Figure 1) 7-day *in-vivo* treatment by MLi-2 significantly reduced phospho-LRRK2 (pLRRK2) and phospho-Rabs in VKI.** A: Representative blots of LRRK2 and Rab protein phosphorylation. MLi-2 treatment significantly reduced pS935-LRRK2 in all genotypes (B: N=4 for all groups; Two-way ANOVA followed by Sidak’s multiple comparison: Genotype: F_2, 18_ =2.69, p =0.09; Treatment: F_1, 18_=157.9, p<0.0001 Genotype x Treatment: F_2, 18_ = 2.35, p=0.12.) and downstream Rab10, 12 and 29 phosphorylation with a clear interaction between genotype and treatment. (**C-E**: N=4 for all groups; Two-way ANOVA followed by Sidak’s multiple comparison: **C**: Genotype: F_2, 18_ =33.48, p<0.0001; Treatment: F_1, 18_=68.81, p<0.0001 Genotype x Treatment: F_2, 18_ = 10.69, p=0.0009; **D**: Genotype: F_2, 18_ =43.29, p<0.0001; Treatment: F_1, 18_=479.3, p<0.0001 Genotype x Treatment: F_2, 18_ = 15.96, p=0.0001; **E**: Genotype: F_2, 18_ =20.89, p<0.0001; Treatment: F_1, 18_=64.69, p<0.0001 Genotype x Treatment: F_2, 18_ = 15.95, p=0.0001). *p < 0.05, **p < 0.01, ***p < 0.001, ****p < 0.0001. Error bars are ±SEM.s

**Figure S2: (Related to figure 6) Dopaminergic markers in mice following 7-day MLi-2 and AMPH induced hyperlocomotion. A:** Representative blots of dopaminergic markers. **B:** Vesicular monoamine transporter (VMAT2) levels are comparable among WT, VKI and Haplo animals following MLi-2 and AMPH. (N=4 animals per group; Two-way ANOVA: Genotype: F _(2, 18)_ = 2.30, p=0.13; Treatment F_1, 18_=4.67; p<0.05; Interaction F_2, 18_= 0.53, p=0.60) **C:** Vehicle treated Haplo animals exhibit significant reduction in phosphor-serine40 TH levels compared to its WT counter part (N=4 animals per group; Two-way ANOVA: Genotype: F _(2, 18)_ = 3.64, p<0.05; Treatment F_1, 18_=0.33; p=0.57; Interaction F_2, 18_= 1.12, p=0.34; Šídák’s multiple comparisons test: WT vehicle vs. Haplo vehicle p<0.05). *p < 0.05, **p < 0.01, ***p < 0.001. Error bars are ±SEM.s

## Acknowledgement

The project is funded by the Lee and Lauren Fixel Chair in Parkinson’s research (M.J.F). Canadian Institutes of Health Research (CIHR) Operating Grant (A.J.M., M.J.F), Weston foundation (M.J.F) and NIH grant R00 R00NS110878 (M.S.M).

## Author contribution

M.B.: conceptualization and project design, data curation, formal analysis, writing original draft, review and editing J.F.: conceptualization and project design, data curation and formal analysis, review and editing, I.T.: conceptualization and project design, data curation, formal analysis, review and editing, S.W.: data curation, formal analysis, review and editing. D.G.: data curation, formal analysis, review and editing. I.D.: data curation, formal analysis, review and editing. G.W.: data curation, formal analysis, review and editing. A.M: Resources, training, and supervision, review and editing. M.S.M.: Resources, training, and supervision, review and editing. H.K.: Resources, training and supervision, review and editing. M.J.F.: Resources, conceptualization and project design, training and supervision, funding acquisition, critical review, interpretation and editing.

