## Supplementary figures and images for "Inhibition of LRRK2 kinase activity rescues deficits in striatal dopamine dynamics in VPS35 p.D620N knock-in mice"

### Supplemental Figure 1

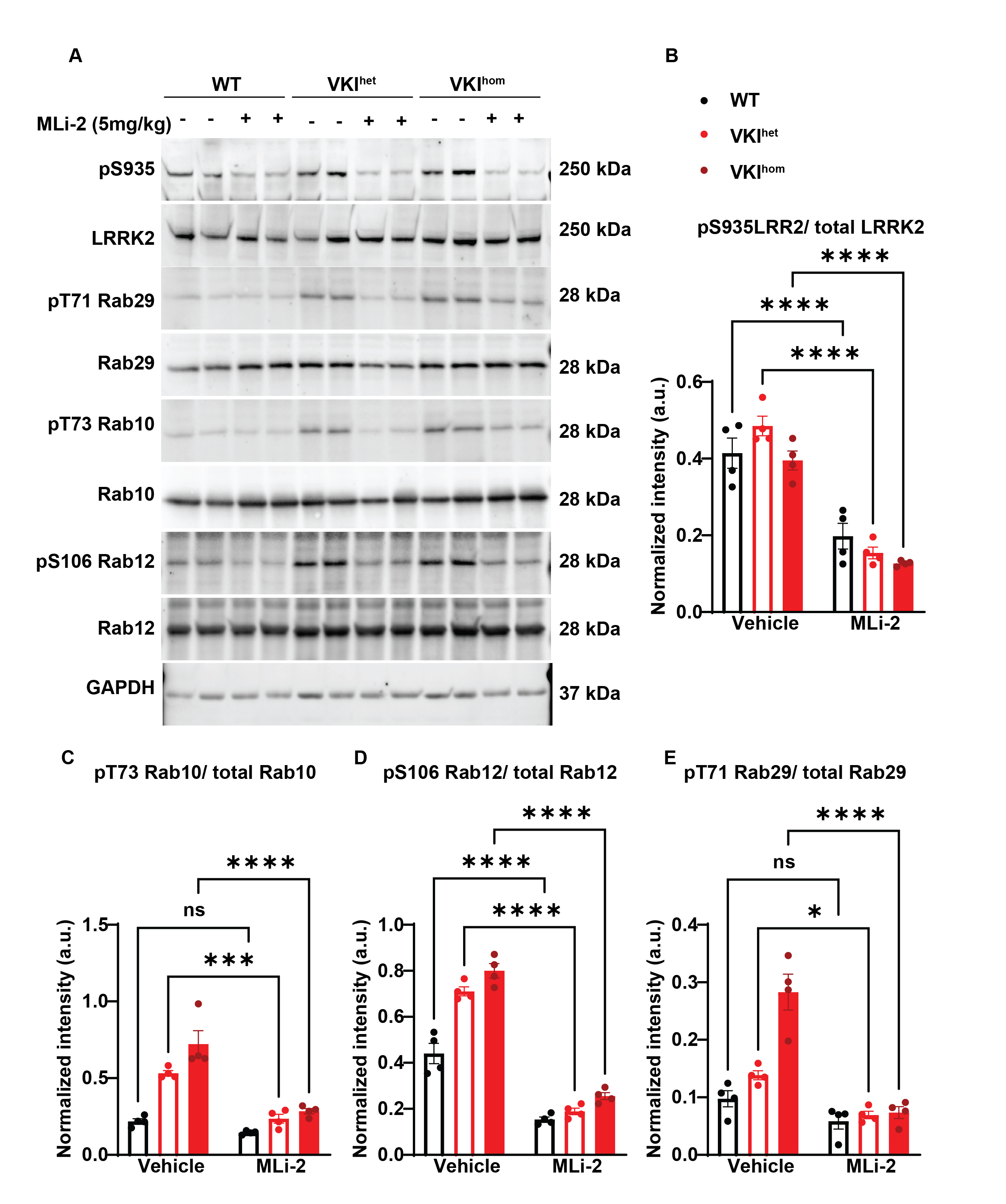

### Supplemental Figure 2

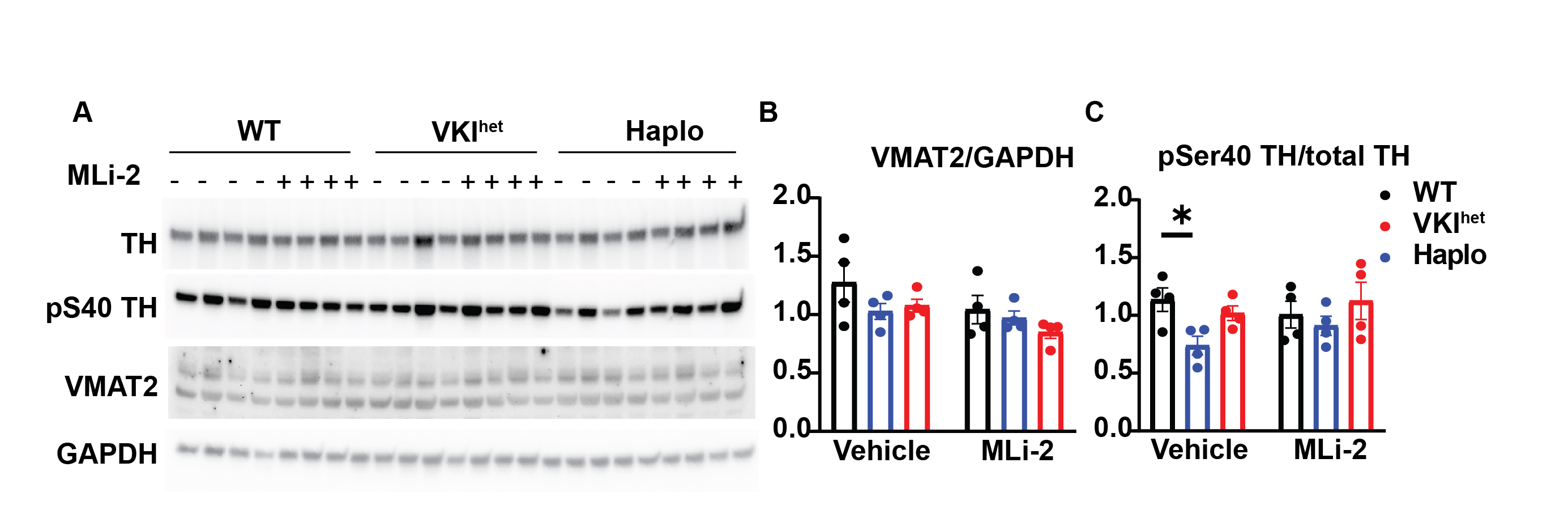
