## Supplemental Table 1 for "Inhibition of LRRK2 kinase activity rescues deficits in striatal dopamine dynamics in VPS35 p.D620N knock-in mice"

Table S1: List of antibodies and dilutions.

| primary antibody | source | host species | Monoclonal/ Polyclonal | catalog number | primary concentration | secondary concentration |
| --- | --- | --- | --- | --- | --- | --- |
| VPS35 | Abnova | Mouse | Monoclonal | H00055737-M02 | 1:2000 | 1:2000 |
| DAT | Millipore | Rat | Monoclonal | MAB369 | 1:1000 | 1:1000 |
| TH | Millipore | Rabbit | Polyclonal | ab152 | 1:1000 | 1:1000 |
| GAPDH | Thermo Scientific | Mouse | Monoclonal | MA5-15738 | 1:2000 | 1:2000 |
| [MJF-R21] RAB10 (phospho T73) | Abcam | Rabbit | Monoclonal | ab230261 | 1:1000 | 1:1000 |
| RAB10 | Abcam | Rabbit | Monoclonal | ab237703 | 1:1000 | 1:1000 |
| Rab12 pS106 (MJF-R25-9) | Abcam | Rabbit | Monoclonal | ab256487 | 1:1000 | 1:1000 |
| RAB 12 | Proteintech | Rabbit | Polyclonal | 18843-1-AP | 1:1000 | 1:1000 |
| RAB29 (phospho T71) [MJF-R24-17-1] | Abcam | Rabbit | Polyclonal | ab241062 | 1:1000 | 1:1000 |
| RAB29 [MJF-R30-124] | Abcam | Rabbit | Polyclonal | ab256526 | 1:1000 | 1:1000 |
| LAMP1 | Abcam | Rabbit | Monoclonal | ab208943 | 1:1000 | 1:1000 |
| SQSTM1 (P62) | Abcam | Rabbit | Polyclonal | ab109012 | 1:1000 | 1:1000 |
| LRRK2/Dardarin (IgG2a) | NeuroMab | Mouse | Monoclonal | N241A/34 | 1:500 | 1:500 |
| LRRK2 p-S935 | Abcam | Rabbit | Monoclonal | ab133450 | 1:500 | 1:500 |
| VMAT2 | Gift from lab of Dr. Gary Miller | Rabbit | Antisera | - | 1 :2000 | 1:2000 |
| mouse anti-rabbit IgG-HRP | Santa Cruz | Mouse | Polyclonal | sc-2357 |  |  |
| anti-mouse IgG-HRP | Santa Cruz |  | Polyclonal | [sc-516102](http://www.scbt.com/product.php?datasheet=2314) |  |  |
| goat anti-rat IgG | biolegend | Goat | Polyclonal | poly4054 |  |  |
